# Spectrum Degradation of Hippocampal LFP During Euthanasia

**DOI:** 10.1101/2020.12.28.424611

**Authors:** Y. Zhou, A. Sheremet, J. P. Kennedy, Nicholas M. DiCola, Carolina B. Maciel, Sara N. Burke, A.P. Maurer

## Abstract

The hippocampal local field potential (LFP) exhibits a strong correlation with behavior. During rest, the theta rhythm is not prominent, but during active behavior, there are strong rhythms in the theta, theta harmonics, and gamma ranges. With increasing running velocity, theta, theta harmonics and gamma increase in power and in cross-frequency coupling, suggesting that neural entrainment is a direct consequence of the total excitatory input. While it is common to study the parametric range between the LFP and its complementing power spectra between deep rest and epochs of high running velocity, it is also possible to explore how the spectra degrades as the energy is completely quenched from the system. Specifically, it is unknown whether the 1/f slope is preserved as synaptic activity becomes diminished, as low frequencies are generated by large pools of neurons while higher frequencies comprise the activity of more local neuronal populations. To test this hypothesis, we examined rat LFPs recorded from the hippocampus and entorhinal cortex during barbiturate overdose euthanasia. Within the hippocampus, the initial stage entailed a quasi-stationary stage when the LFP spectrum exhibited power-law feature while the frequency components over 20 Hz exhibited a power decay with a similar decay rate. This stage was followed by a rapid collapse of power spectrum towards the absolute electrothermal noise background. As the collapse of activity occurred later in hippocampus compared with medial entorhinal cortex or visual cortex, it suggests that the ability of a neural network to maintain the 1/f slope with decreasing energy is a function of general connectivity. Broadly, these data support the energy cascade theory where there is a cascade of energy from large cortical populations into smaller loops, such as those that supports the higher frequency gamma rhythm. As energy is pulled from the system, neural entrainment at gamma frequency (and higher) decline first. The larger loops, comprising a larger population, are fault-tolerant to a point capable of maintaining their activity before a final collapse.

## 1. Introduction

For over five decades it has been evident that the hippocampal local-field potential (LFP) activity is strongly correlated with behavior [Vanderwolf, 1969, Buzsaki, 2005]. The most prominent feature of hippocampal LFP, the 8-9 Hz theta rhythm, is reported to increase in power with increasing running speed [Whishaw and Vanderwolf, 1973, Morris and Hagan, 1983]. and change its shape from sinusoidal to sawtooth waves [Green and Petsche, 1961, Stumpf, 1965, Buzsaki et al., 1983, Terrazas et al., 2005] associated with the development of high order theta harmonics during faster movement [Leung, 1982, Leung et al., 1982, Sheremet et al., 2016, Zhou et al., 2019]. Apart from theta, the power of the higher frequency gamma rhythm (60-120 Hz) also increases with respect to running speed [Chorbak and Buzsaki, 1998, Chen et al., 2011, Ahmed and Mehta, 2012, Kemere et al., 2013]. The increase in gamma power at faster running speeds is also associated with enhanced thetagamma coupling [Sheremet et al., 2018]. These studies imply that when the rat is in an active behaving state, the hippocampus receives strong barrages of synaptic input [Shu et al., 2003], giving rise to an organization of activity across all spatial and temporal scales [Sheremet et al., 2020].

The hippocampal LFP is often decomposed through Fourier analysis, providing the power spectral density. The power spectra density takes the form of 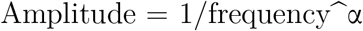, where alpha falls is a value between 0 and two (t characteristic of a pink noise spectra; Buzsaki, 2006). While this decomposition treats different frequency bands as independent signals, the theta and gamma oscillations are not isolated rhythms. In addition to being coupled to each other, theta and gamma also exist against a “background” of activity. While the exact nature of this background is not well understood, it is often attributed to either the result of interacting oscillators [Buzsaki, 2006] or the consequence of a broadband, arrhythmic activity that is distinct from oscillations [Hesse and Thilo, 2014]. Importantly, similar to theta and gamma, the slope of the spectra that is attributed to the background has also been found to change with increasing running speed [Sheremet et al., 2019b], suggesting that it is sensitive to increased synaptic currents and may be interrelated with the rhythms that are often ascribed as being independent from the background spectra.

One explanation for why power is lower at higher frequencies is the energy cascade across multiple oscillators. Specifically, axonal conduction delay and synaptic time constants determine the frequency in which the populations of neurons can rhythmically engage. Faster oscillations require rapid communication, suggesting that a rhythm like gamma would be local, generated by a small population of neurons. The organization of a larger pool of neurons is rate limited by communication. Therefore, the propagation of activity between the hippocampus and cortex, will take the form of reciprocal volleys or traveling waves at a slower frequency. But with the increased engagement of the neuronal population, an oscillation such as theta will be one order of magnitude larger [Buzsaki and Draguhn, 2004]. Importantly, these oscillators – among many others-are not independent but one in the same [Bullock et al., 1995, Sheremet et al., 2019a].As these rhythmic processes are intertwined, activity across the system will generate a spatio-temporal interaction that gives rise to the 1/f “pink” spectra [Buzsaki, 2006]. However, other theories regarding the organization of the power spectra suggest that there are two independent biological processes that support aperiodic, noise (the 1/f slope) and the rhythmic frequencies, the “peaks” above the slope [He, 2014, Donoghue et al., 2020]. This difference in interpretation of the slope may be related to the common practice of discarding or compensating/”whitening” out the slope [Buzsaki, 2006, Donoghue et al., 2020]. The 1/f slope is more often ignored or considered to be “arhyth-mic” activity in the absence of biological justification. That is, the underlying mechanisms that support the 1/f slope in the spectra should be appreciated and well-understood before either being carved or discounted.

The 1/f slope of the spectra is a consistent feature of the LFP and has relationships to the canonically defined rhythms theta and gamma. For instance, both theta and gamma increase in power and frequency with running speed [Ahmed and Mehta, 2012, Sheremet et al., 2019a] which co-occurs with a change in the 1/f slope [Sheremet et al., 2019b]. However, the 1/f slope is present during sleep and quiescent periods, when the power in the theta and gamma bands is at the 1/f slope level (for instance, see figure 4 Buzsáki et al., 2003). This suggests that the 1/f slope is a persistent and robust feature of neural activity. That is, the 1/f slope may be supported by large scale neural activity necessary to maintain homeostasis, and manipulating the spectra require approaches that alter a large swath of synaptic inputs, such as localized cooling [Petersen and Buzsáki, 2020], focal brain lesion [Mitchell et al., 1982, Buzsaki et al., 1987, Bragin et al., 1995, Fyhn et al., 2004], and study the dying brain. In the current study, we investigated the spectra against the background of euthanasia by barbiturate overdose, in which the spectra degrades to a complete collapse.

Prior studies on LFP changes with euthanasia have observed a surge of global and highly coherent gamma oscillation after cardiac arrest [Borjigin et al., 2013].This has led to studies exploring hippocampal physiology in the context of near-death experience [Parnia and Fenwick, 2002], theoretically supported by bursts of high-frequency activity [Zhang et al., 2019]. While near-death experience is interesting, this topic is beyond the scope of the present study. Rather, we explored the changes from the perspective of the energy cascade hypothesis. If slow frequency perturbations provide the energy to drive higher frequency oscillations, then we predicted that when the system is significantly challenged, the high frequency rhythms and the 1/f slope will be the first to be compromised. Low frequency activities, being generated by larger more distributed populations, will be more robust, maintaining power for an extended period of time. In accord with this, we initially observed high-frequency events, similar to other publications [Zhang et al., 2019], followed by a steepening of the 1/f carried by a recession of high frequency power. At end stage, the LFP was mainly characterized by low frequency, high amplitude activity prior to the final collapse of all activity. This phenomenon was observed across the hippocampus and entorhinal cortex, although the spectral integrity persisted in the hippocampus for relatively longer.

## 2. Materials and Methods

### 2.1. Subjects and Behavioral Training

All behavioral procedures were performed following protocols approved by the Institutional Animal Care and Use Committee at the University of Florida as well as those set forth by the National Institute of Health. The present study consisted of 3 male hybrid Fisher344-Brown Norway rats (Taconic) ranging from 4 to 10 months of age (r730, r782, and r1074 all male). Animals were singly housed and allowed to acclimate for one week after arrival. The colony room maintained a reversed 12-12 hour light-dark cycle with all behavior taking place during the dark period. Behavioral shaping began with training animals to run counterclockwise on a circular track one meter in diameter for a food reward (pieces of cereal marshmallow, Medley Hills Farm, Ohio). During this time, the animal’s weights were slowly reduced to 85% of their ad lib. weight. Once a criterion of at least 30 laps in 15 minutes was reached, animals were implanted unilaterally with silicon probes. One probe was implanted in the dorsal hippocampus (HPC) in all the three rats. As to rat 730 and rat 782, another probe was implanted in the medial entorhinal cortex (MEC). The probes used for animals that received implants in the HPC and MEC were custom single shank, 32 channel probes (NeuroNexus; Ann Arbor, MI) with an area of 177 μm2 and a site spacing of 60 μm. The animal where only the HPC was targeted received single shank 64 channel probe (L3 series; Cambridge NeuroTech; Cambridge, UK) with an area of 165 μm2 and a site spacing of 50 μm. Prior to surgery, all probes were cleaned by soaking in a solution of 7% detergent (Contrad 70 Liquid Detergent; Decon Labs; King of Prussia, PA) in deionized water followed by rinsing with deionized water.

### 2.2. Surgical Procedures

All surgical procedures were performed following protocols approved by the Institutional Animal Care and Use Committee at the University of Florida as well as those set forth by the National Institute of Health. Animals were placed in an induction chamber and sedated with 3-5% Isoflurane. After loss of muscle tone, they were moved to a nose cone and the top portion of the head was carefully shaved to avoiding cutting any whiskers. Next, the animal was transferred to the nose cone of the stereotaxic frame, where the head was gently secured with using ear and incisor bars. During this portion and for the remainder of the procedure, anesthesia was maintained using an Isoflurane dose between 1% and 2.5% while periodically monitoring respiration. Body temperature was maintained using an electric heating pad with feedback via rectal thermometer. The eyes were protected by applying ophthalmic ointment and shielding from direct light. Prior to the initial incision, the top of the head was cleaned using several cycles of povidone-Iodine and alcohol. An incision was made starting just behind the eyes and continuing to the back of the skull. The skin was retracted, and blunt dissection was used to expose the surface of the skull. Bleeding was managed using a cautery pen (Bovie Medical; Clearwater, FL). After thoroughly cleaning the skull, measurements from a stereotaxic arm were used to ensure that the skull was leveled. Next, bregma and the electrode implant locations were marked on the skull with the cautery pen for visual reference. A total of seven anchor screws were placed into the skull to serve as attachment points for the headcap. One screw over the cerebellum and one screw over the cortex were attached to wires that would serve as the reference and ground locations, respectively. A small amount of luting cement (C&B Metabond; Parkell Inc; Edgewood, NY) was applied to the screws to provide a foundation for the rest of the headcap. Care was taken to avoid covering bregma and the implant sites. Craniotomies were drilled at the implant sites and the dura was removed, taking care to not damage the cortex. Bleeding was managed using saline irrigation or sterile gauze. Probes targeting the dorsal HPC were implanted at −3.2 mm AP; 1.5 mm ML to bregma; −3.7 mm DV to dura. Coordinates targeting the MEC were −0.5 mm AP to the transverse sinus, 4.6 mm ML to bregma, angled 30 degrees posteriorly, and −5.78 mm DV to dura. After implantation, the craniotomies were sealed with a surgical silicone adhesive (Kwik-Sil; World Precision Instruments; Sarasota, FL). Dental acrylic (Grip Cement, 675571 (powder) & 675572 (solvent); Dentsply Caulk; Milford, DE) was then applied to secure the probes and connectors in place. The ground and reference wires were soldered to the appropriate wires on the probe connectors and the reference wire was isolated using dental acrylic. Lastly, copper mesh was shaped into a small bowl around the headcap to serve as physical protection and secured with dental acrylic. The ground wires were soldered to the copper mesh to minimize the danger of electrostatic discharge. Immediately following the removal of the anesthetic, 10 ml of sterile saline and a dose of 1.0 mg/kg meloxicam (Boehringer Ingelheim Vetmedica, Inc; St. Joseph, MO) were administered subcutaneously. The animals were placed on a heating pad and monitored until fully mobile and capable of eating. Post-surgical care included a second dose of meloxicam 24 hours later as well as 5ml of oral antibiotics (40 mg/ml Sulfamethoxazole & 8mg/ml Trimethoprim Oral Suspension; Aurobindo Pharma Inc; Dayton, NJ) mixed into their food for seven days. Animals were monitored for one week following surgery to ensure no physical or behavioral abnormalities were observed before testing began.

### 2.3. Euthanasia Electrophysiology

After completing all other behavioral experiments, animals were recorded from for 15 minutes in the usual resting container to establish a baseline for the LFP data and then received a lethal dose of SomnaSol (390 mg/ml pentobarbital sodium, 50 mg/ml phenytoin sodium; Henry Schein; Melville, NY) injected intraperitoneally. LFP recording continued throughout the injection and for 10 to 15 minutes after the animal no longer exhibited a nociceptive withdrawal reflex. The animal was then immediately perfused with 4% paraformaldehyde, and the brain extracted and prepared for histology to verify electrode locations.

### 2.4. Data process and spectral analysis

The LFP data were analyzed in MATLAB (The MathWorks, Natick, MA) using custom-written code as well as code imported from the HOSAtoolbox [Swami et al., 2000]. Raw LFP records sampled at 24 kHz (Tucker-Davis system) were low-pass filtered down to 1 kHz. The spectrogram were calculated based on discrete Fourier transform with window length of 1 second and 50% overlap. The power correlograms for each evolution stage were obtained by estimating the correlation coefficients between all the frequency pairs in the result of spectrogram. The power spectra during the euthanasia were estimated for every 100 seconds. Within each LFP interval, the power spectrum was obtained via the standard Welch’s method [Welch, 1967] with window length of 1 second and 50% overlap. The power law exponential was obtained by linearly fit the log-log power spectrum in the frequency range from 20 Hz to 80 Hz.

The coherence between two time series is the modulus of their cross-spectrum normalized by their power spectra. The cross-spectrum and power spectra were estimated with window length of 1 second and 50% overlap. To demonstrate the time evolution of coherence, a sliding window with a length of 20 seconds and a step increment of 5 seconds was applied. In each window, the coherence between two time series in frequency range from 2 Hz to 500 Hz were estimated.

The asymmetry and skewness of LFP were obtained from the bispectral analysis with a window length of 1 second and 50% overlap. The bispectrum has been thoroughly reviewed in terms of both statistical and mathematical background [Harris, 1967] as well as its application to nonlinear wave interaction [Kim and Powers, 1979]. In the field of neuroscience, bispectral analysis (the Fourier transform of the third-order cumulant) was used to quantify the degree of phase coupling between the frequencies of the LFP, whereas the bicoherence quantifies the degree of cross-frequency coupling independent from the amplitude [Barnett et al., 1971, Ning and Bronzino, 1989, Sigl and Chamoun, 1994, Bullock et al., 1997, Muthuswamy et al., 1999, Hagihira et al., 2001, Li et al., 2009, Wang et al., 2017, Avarvand et al., 2018]. The crossbispectrum analysis is similar with the bispectrum analysis but the frequency components are from two time series [Lii and Helland, 1981]. In our study, as the stage length differs across regions, the cross-bicoherence between HPC and MEC were estimated over the shorter stage length.

The decay rate of each frequency component was derived from the power time series. Power time series of frequency component *ω* was defined as the variance of filtered LFP in the frequency band *[ω* — 0.5Hz, *ω* + 0.5Hz]. The power time series was either calculated with a window length of 1 seconds and a step increment of 0.5 second (figure 3.3 A), or with a window length of 10 seconds and a step increment of 2 second (figure 3.5 C). The decay/grow constant was defined as the ratio of the time derivative of the power time series to the power time series itself. The obtained result was average over the time window of 20 seconds to eliminate fast power oscillations, and was plotted every 5 seconds. The transition period from the first to the second stage is defined as the moment with the fastest averaged decay rate over 100 Hz. The transition period from the second to the third stage is defined as the moment with the fastest averaged decay rate from 4 to 60 Hz.

**Figure 3.1.**
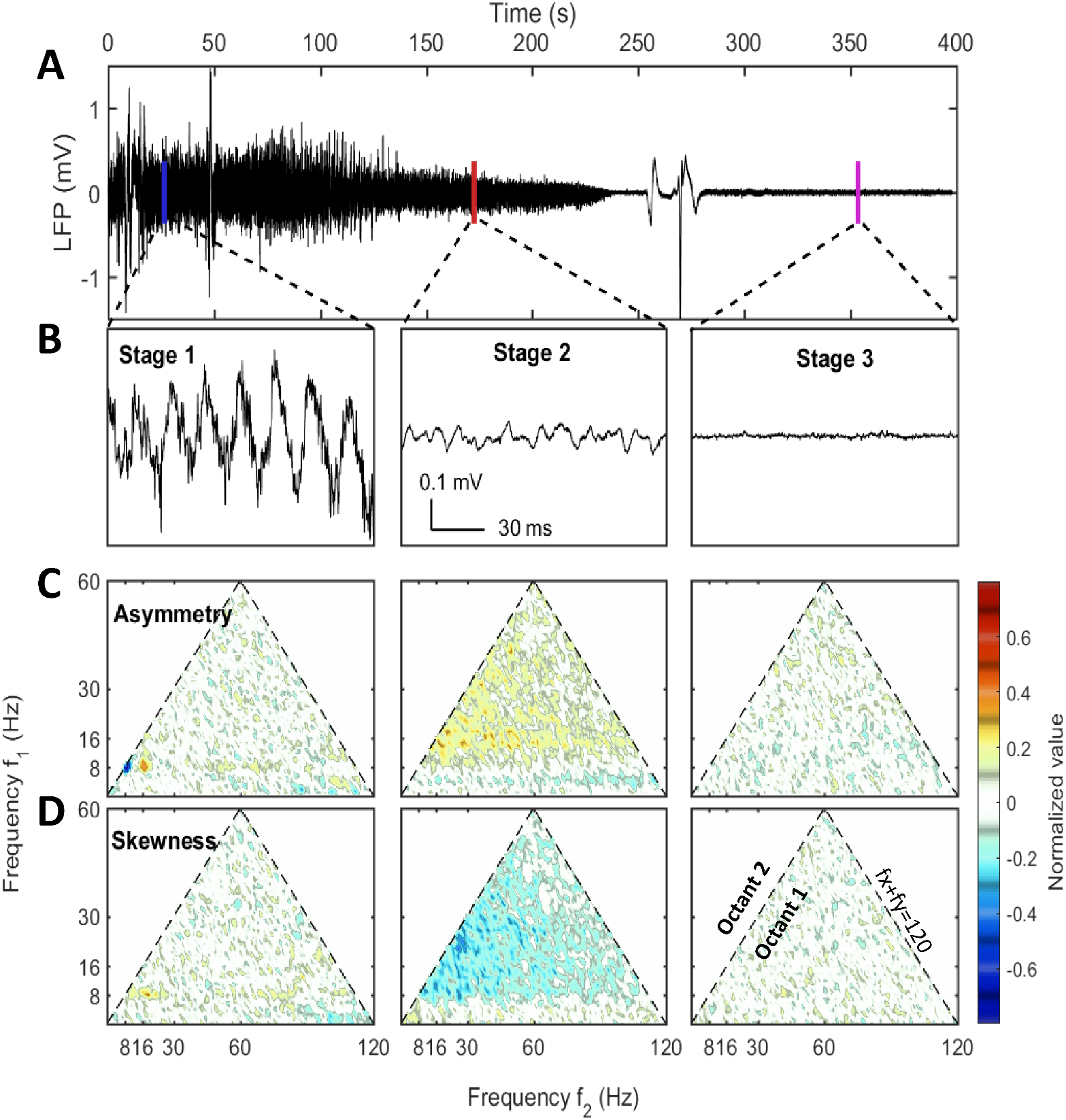
LFP examples at different stages and their asymmetry and skewness. A) The raw LFP trace recorded in CA1 pyramidal layer after the injection (*t* = 0) which showed the overall neural activity evolution during euthanasia. B) LFP samples of 1 second selected from three stages. The corresponding windows were indicated in panel A. C) & D) The asymmetry and the skewness of LFP at three stages obtained from bispectral analysis [Haubrich and MacKenzie, 1965, Masuda and Kuo, 1981, Sheremet et al., 2016]. Both the horizontal axis and the vertical axis represented frequency. Each point in the plot was the asymmetry or skewness estimated for the frequency triad (*f_x_,f_y_,f_x_ +f_y_*). Only the first octant (*f_x_* ≥ 0, *f_y_* ≥ 0, *f_x_* ≥ *f_y_*) is presented as it contained all the non-redundant information due to the symmetry of bispectrum. The first octant is bounded by the dashed line *+f_y_* = 120 Hz indicating the upper bound of frequency range. An oscillation has asymmetry if the wave peak or trough doesn’t stand at the center of adjacent zero-across points. An oscillation has skewness if the wave height distribution is not symmetric about its mean value. Both asymmetry and skewness indicate the system is nonlinear. Data from rat 782.

**Figure 3.2.**
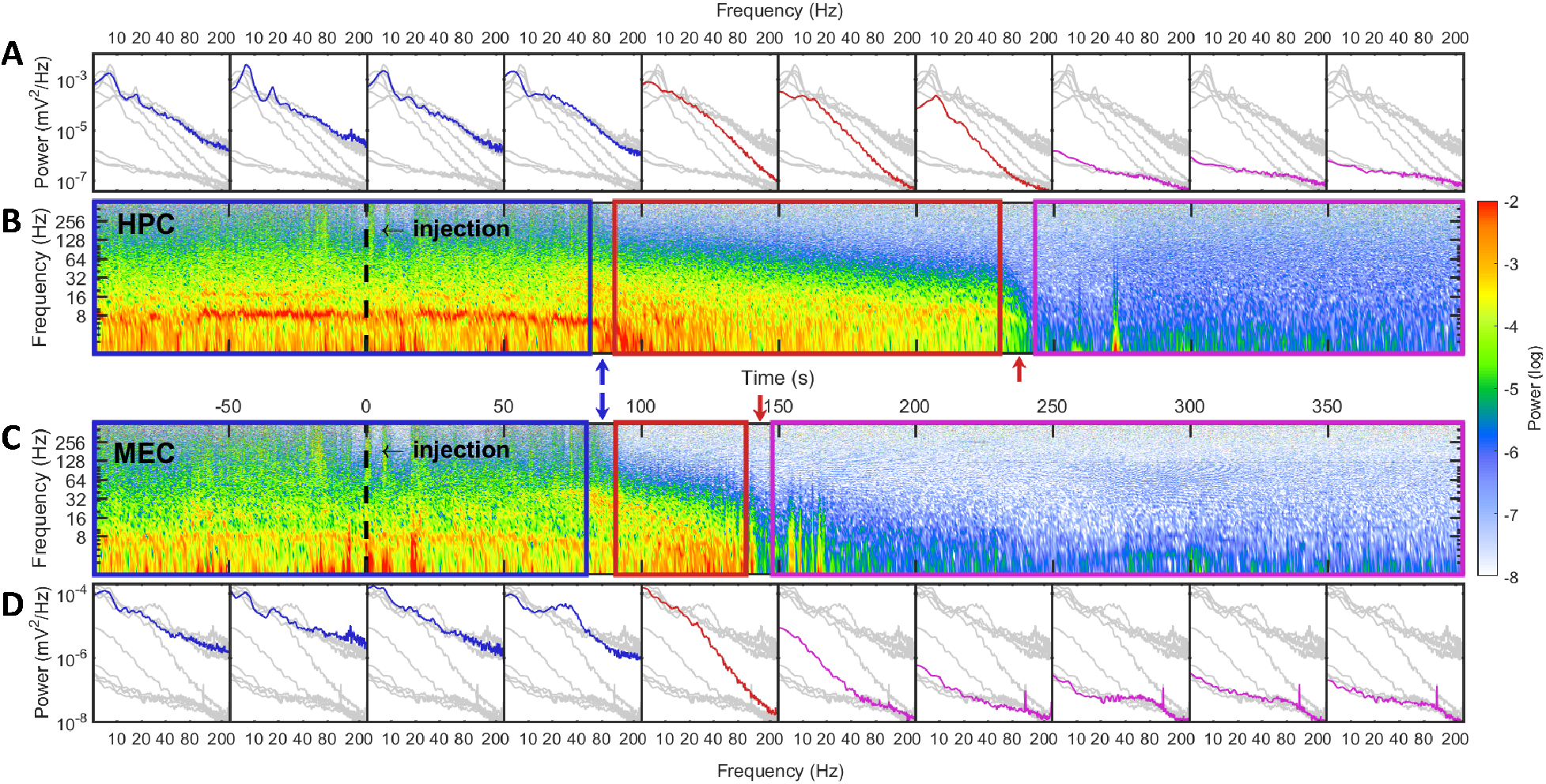
Spectrum degradation in HPC (top two rows) and MEC (bottom two rows). LFPs in HPC and MEC were collected simultaneously from two probes. A) The development of spectra estimated every 50 seconds. Spectra estimated within other time intervals were indicated as gray lines for comparison. B) The spectrogram of hippocampal LFP where the injection time was marked as a dashed line at 0 second. The power in spectrogram was normalized by the maximum power during the euthanasia process. Based on the spectrogram and the development of power spectrum, there were three stages during euthanasia: stage 1 was the pre-effective stage which included the pre-injection period and a short period after injection. In spectrogram it was marked with a blue box and in spectra they were blue lines; stage 2 was the quasi-stationary decay stage marked as a red box in spectrogram and red lines in spectra; stage 3 was the quasi-white noise stage marked as a magenta box in spectrogram and magenta lines in spectra. Between these stages there were two rapid transition periods marked with blue and red arrows. C) The spectrogram of LFP recorded in MEC. D) The development of spectra in MEC. Similar with the observation in HPC, three spectrum evolution stages and two transition periods can be observed. The transitions from stage 1 to stage 2 occurred almost simultaneously in HPC and MEC. However, MEC had a much shorter stage 2 and the spectra collapsed to an almost white noise spectrum quickly. Data from rat 782.

**Figure 3.3.**
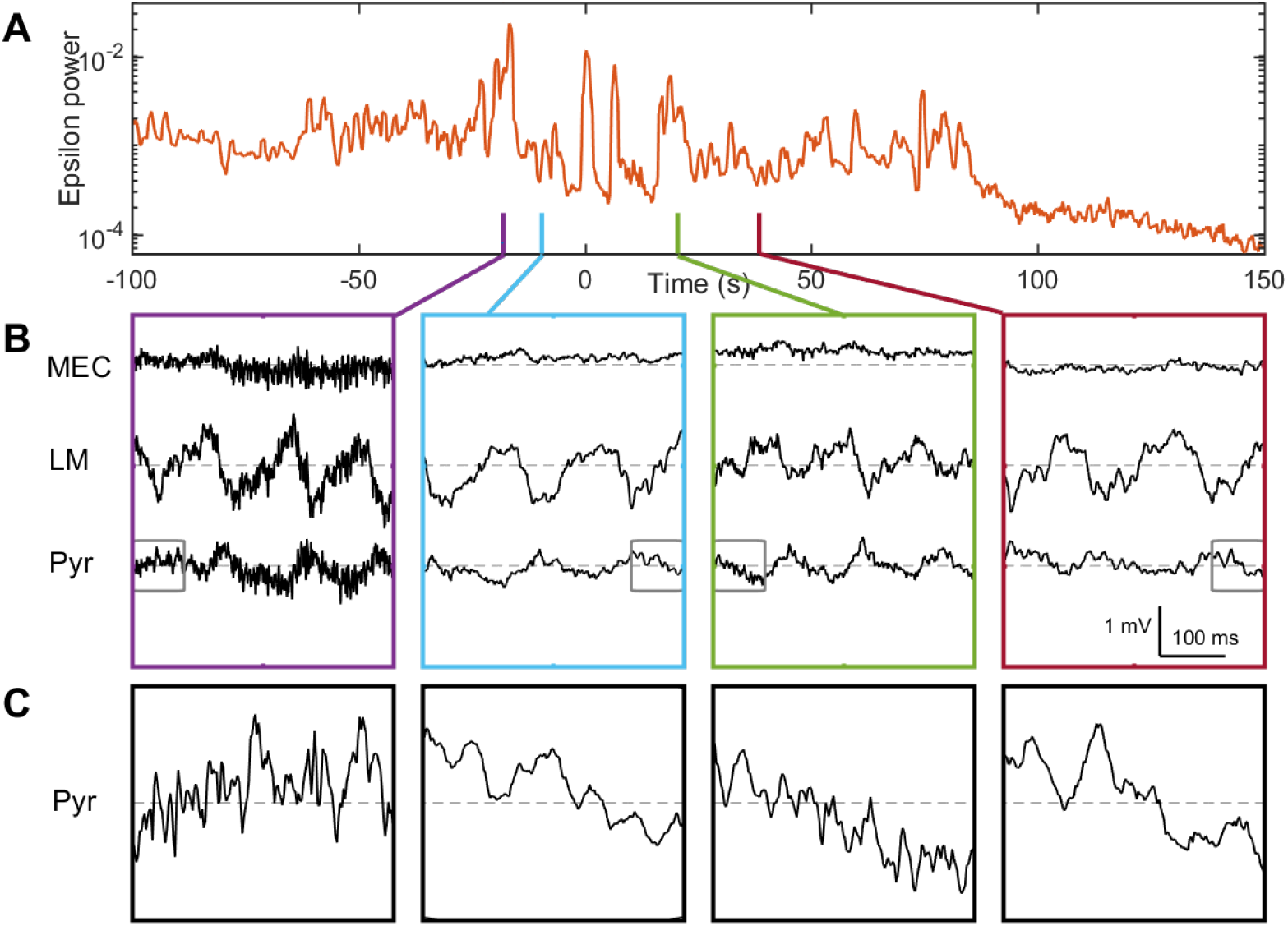
Non-stationary hippocampal LFP during stage 1. A) >100 Hz power evolution estimated with window length of 1 sec and step increment of 0.5 sec. Stage 1 approximately ended at *t* = 80 sec. Part of stage 2 was included for comparison. Four instances were marked on time axis. Instances marked with purple and green had strong >100 Hz power, while the instances marked with blue and red had weak >100 Hz power. B) Raw LFPs recorded in MEC, CA1 lacunosum-moleculare (LM), and CA1 pyramidal layer (Pyr) of four instances marked in panel A. C) Zoomed-in LFPs in CA1.Pyr indicated by gray boxes in panel B. Embedded high frequency oscillations were stronger during instances with high >100 Hz power.

## 3. Results

### 3.1. Three Stages during Spectrum Degradation

After the injection of SomnaSol, LFP exhibited three distinguishable stages with two rapid transition periods between these stages in all the animals.

#### 3.1.1. Pre-effective stage

The first stage (marked as blue box in figure 3.2) was described as the pre-effective stage,or stage 1. This stage was indistinguishable from baseline (pre-injection). Thus in figures 3.1& 3.5, both the pre-injection (t<0) and the pre-effective stages were marked as stage 1. In this stage, LFP was dependent on behaving state of the rat where there were strong theta and gamma rhythms at high running velocity [Whishaw and Vanderwolf, 1973, Morris and Hagan, 1983, Chen et al., 2011, Ahmed and Mehta, 2012, Kemere et al., 2013, Zheng et al., 2015, Sheremet et al., 2019a]. A sample of hippocampal LFP from stage 1 (figure 3.1B) reveals that theta can express significant deviations from a sinusoid [Buzsaki et al., 1983, Terrazas et al., 2005]. This nonsinusoidal waveform is related to high order theta harmonics due to the nonlinearity of hippocampal LFP [Scheffer-Teixeira and Tort, 2016, Sheremet et al., 2016, Zhou et al., 2019]. In statistical analysis, the lowest order nonlinear character of the system can be described by bispectrum [Hasselmann et al., 1963]. The real and imaginary part of the bispectrum characterizes the skewness (an example being a cnoidal wave) and the asymmetry (“sawtooth” shaped wave) of the distribution. The sawtooth aspect of the theta wave, with steep wave front (from trough to peak), corresponded to the negative asymmetry (figure 3.1 C) at the frequency triad (8,8,16) Hz (8 Hz at x-axis, 8 Hz at y-axis and their sum 16 Hz). Apart from that strong negative asymmetry region, frequency triad (8,16,24) Hz exhibited strong positive asymmetry and supported the existence of third order theta harmonic [Schomburg et al., 2014, Zhou et al., 2019, Cowen et al., 2020].

In the spectrogram of hippocampal LFP (figure 3.1 B) during stage 1, there was a strong theta oscillation along with intermittent but distinguishable 2nd and 3rd harmonics. Apart from high power activities in theta and its harmonics bands, intermittent high frequency events can be identified by looking at spectrogram at frequency range over 100 Hz (perhaps analogous to *epsilon* described by others; Canolty et al., 2006, Freeman, 2007, Sullivan et al., 2011, Belluscio et al., 2012). The intermittency was supported by the power evolution (figure 3.3 A) where the >100 Hz power had strong variation in the log-scale plot. Comparing spectrogram between LFPs in HPC and MEC(figure 3.1, 5.1 & 5.2, panel B&C), we further discovered strong >100 Hz epochs were synchronized across three brain regions (HPC and MEC), indicating the >100 Hz bursts were not localized events. At the moments with >100 Hz bursts, we can observe strong high frequency oscillations embedded in theta rhythm, and the raw LFP appears “fuzzier” compared with moments with low >100 Hz power (figure 3.3 B&C). Power spectra during stage 1 also supported that the LFPs were dominated by the theta rhythm, and high theta harmonics were prominent in the HPC region (figure 3.2 A&D). At frequency region of >100 Hz, however, power spectrum is not an optimal representation of the process. As we have shown during stage 1, the >100 Hz events were non-stationary their intermittent occurrence contributed to the majority of the power in >100 Hz range (figure 3.4 A). These intermittent structures had a wide frequency support and caused the spectrum to have a slope of −Λļ ≈ −1.5 in log log scale (figure 3.4 B).

**Figure 3.4.**
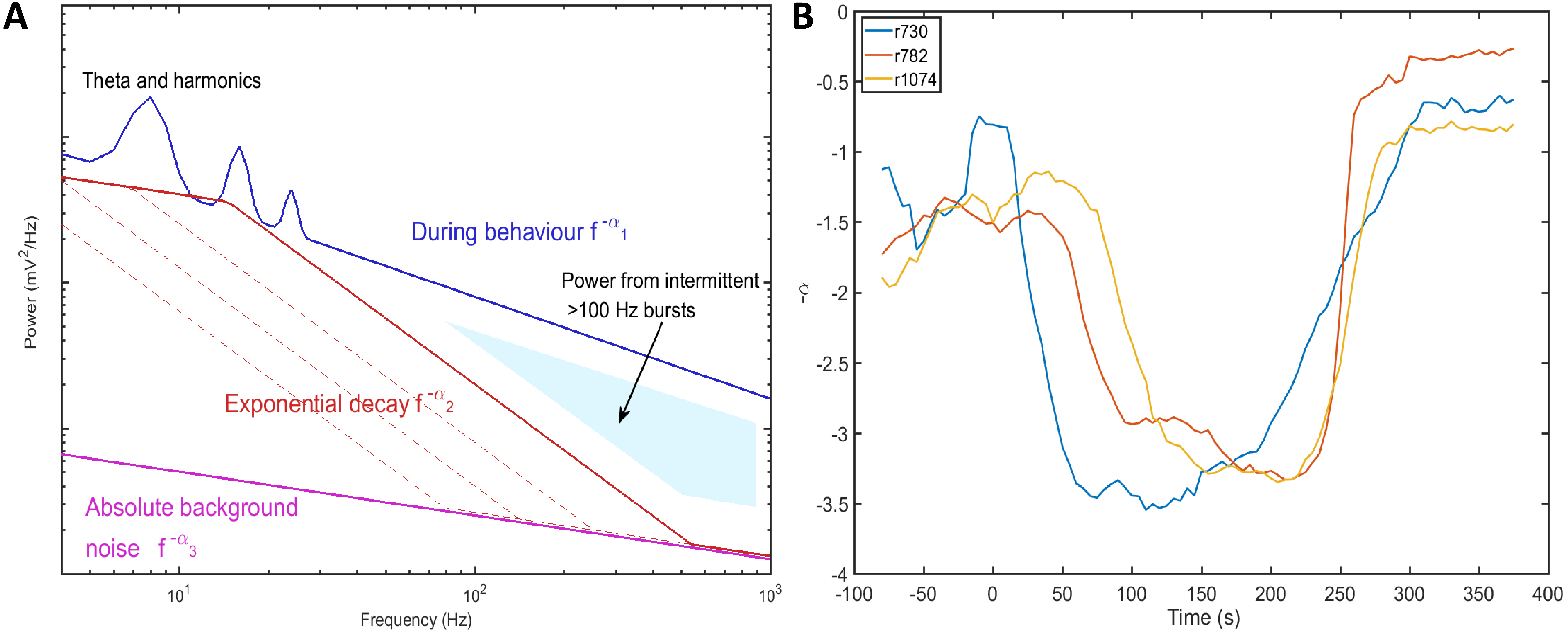
A) The evolution of spectrum during euthanasia. The spectra from the first to the third stages were marked as blue, red and magenta. The power spectra exhibits different slopes across stages. The existence of intermittent epsilon during stage 1 (pre-effective stage) increased the power spectrum at high frequency range and gave rise a relatively flat spectrum. During stage 2 (quasi-stationary decay stage), the spectra evolved as parallel lines (dashed red lines) where the power at high frequency range clung to spectrum at stage 3. During stage 3 (collapse stage), the LFP collapsed and there were limited localized activities, thus the spectrum estimated at this stage was interpreted as absolute background noise which might be impacted by noise from data acquisition instrument. B) Power-law exponents evolution of three rats estimated with window length of 40 seconds and time increment of 5 seconds. The power-law exponent is defined as the slope of power spectrum in log-log plot. Three stages can be identified with *α*_1_ ≈ 1.5, *α*_2_ ≈3.3, and *α*_3_ ≈0.75

#### 3.1.2. Quasi-stationary decay stage

The second stage (marked as red box in figure 3.2) was described as the quasi-stationary decay stage, or stage 2, after the sudden disappearance of theta and >100 Hz bursts (at 82 + 4 seconds after SomnaSol administration). The hippocampal LFP in this quasi-stationary stage exhibited slow power decay over a period of approximately 100 seconds without obvious intermittent structures (figure 3.1 A). Within a narrow time frame, the LFP appeared to be nearly stationary although of lower amplitude compared with stage 1. Traditional 8 Hz theta activity was no longer evident via visual inspection. Rather, the power in theta appears to slow into a lower frequency band (1-4 Hz) which quickly collapses. Interestingly, this surge in lower frequency activity has been reported previously (CITE: https://www.sciencedirect.com/science/article/pii/S030100821930351X). As this lower frequency, 1-4 Hz activity dissipated, a10 Hz dominate oscillation became prominent, characterized by a wide-flat peak and narrow-deep trough (figure 3.1 B). This waveform expressed a negative skewness in the frequency region from 10 to 30 Hz via bispectrum analysis (figure 3.1 D). The spectrogram of LFP during stage 2 was consistent with observations of the time-series. There was low power in >100 Hz region that accompanied the development of a 10 Hz oscillation. In terms of the spectrum evolution, except for the frequency components < 10 Hz, there was a structured power decay over a wide frequency band. Due to the absence of intermittent >100 Hz events, power spectra in stage 2 contained less power in the high frequency range resulting in a steeper slope compared with stage 1 (figure 3.4). The power-law distribution persisted over the entire stage with a slope of — α_2_ ≈ −3.3, but a decaying total variance. This is evident in the straight power contour lines in the spectrogram plot (contour lines were not directly plotted but can be identified from the transition of colors) (figure 3.1 B). As a result, during stage 2 the spectrum experienced a “parallel” evolution that can be interpreted as the entire spectrum having a decay in power along with a shift towards lower frequency. By comparing across different brain regions, we reported such a “parallel” spectrum evolution existed in HPC and MEC(figure 3.2, 5.1 & 5.2). However, although the onsets of stage 2 were synchronized, their lengths varied between the different brain regions. In the HPC region, the power-law spectrum kept evolving after the LFP spectrum in MEC had collapsed to a low power-containing state.

#### 3.1.3. Collapse stage

In the HPC region, the LFP collapsed at 269 + 50 seconds after SomnaSol administration, marking the spectral transition into stage 3, or collapse stage (marked as magenta box in figure 3.2). Oscillation amplitude was small (figure 3.1 B) with occasional large spikes(figure 3.1 A, figure 3.2 B). This oscillation has been described previously as the “wave of death” (WoD; Kaminogo et al., 1998, Van Rijn et al., 2011, Zandt et al., 2011) which is proposed to reflect the massive and simultaneous depolarization of a large number of neurons. This phenomenon most-likely shares a high degree of similarity with cortical spreading depression [Dreier and Reiffurth, 2015] accounting for why the event was not highly correlated between brain regions (figure 3.2 B&C). Pani, Pierpaolo et al. [2018] reported this brain activity can persist for about 120 min after cardiac arrest, because of a slow spreading depolarization caused by irreversible degenerative processes at the cellular level. Apart from the transient spikes, LFP in stage 3 didn’t show significant asymmetry or skewness in the bispectral analysis (figure 3.1 C&D). The power spectrum during stage 3 was flat with a slope of –*α*_3_ ≈ —0.75 and represented the least power containing state of our LFP measurement (figure 3.4). Given the low power level during stage 3, it is possible that the spectrum reflected more of the electro-thermal noise in physical recording environment surrounding the data acquisition system than neural activities.

### 3.2. Global Degradation of Theta and Intermittent >100 Hz Bursts

In the previous section, we stated the stage 1 of spectrum degradation ended with the erosion of observable theta and the disappearance of the intermittent high frequency activity. Comparing spectrogram between HPC and MEC, we discovered that the transition from stage 1 to stage 2 was synchronized across these regions (figure 3.1, 5.1 & 5.2, panel B&C). The spectrogram showed the intermittent high frequency structures vanished after stage 1 (figure 3.2 B&C), and the corresponding power in >100 Hz band decreased rapidly in HPC and MEC (figure 3.5, figure 5.3 & figure 5.4 panel C). The burst suppression could be caused by sodium pentobarbital[Aleman et al., 2015], which enhances postsynaptic responses to the inhibitory actions of GABA or related to hemodynamic collapse and lack of brain perfusion. Although the typical theta oscillation along with its harmonics was no longer visually observable at the end of stage 1 (figure 3.2 B&C), the power in theta band (6-10 Hz) only exhibited a limited decrease from stage 1 to stage 2 in the HPC and MEC regions (figure 3.5, figure 5.3 & figure 5.4 panel C). Therefore, in order to determine if the 6-10 Hz frequency component is a degenerate form of theta or due to a different mechanism, we sought to determine if the band shared features generally associated with theta.

**Figure 3.5.**
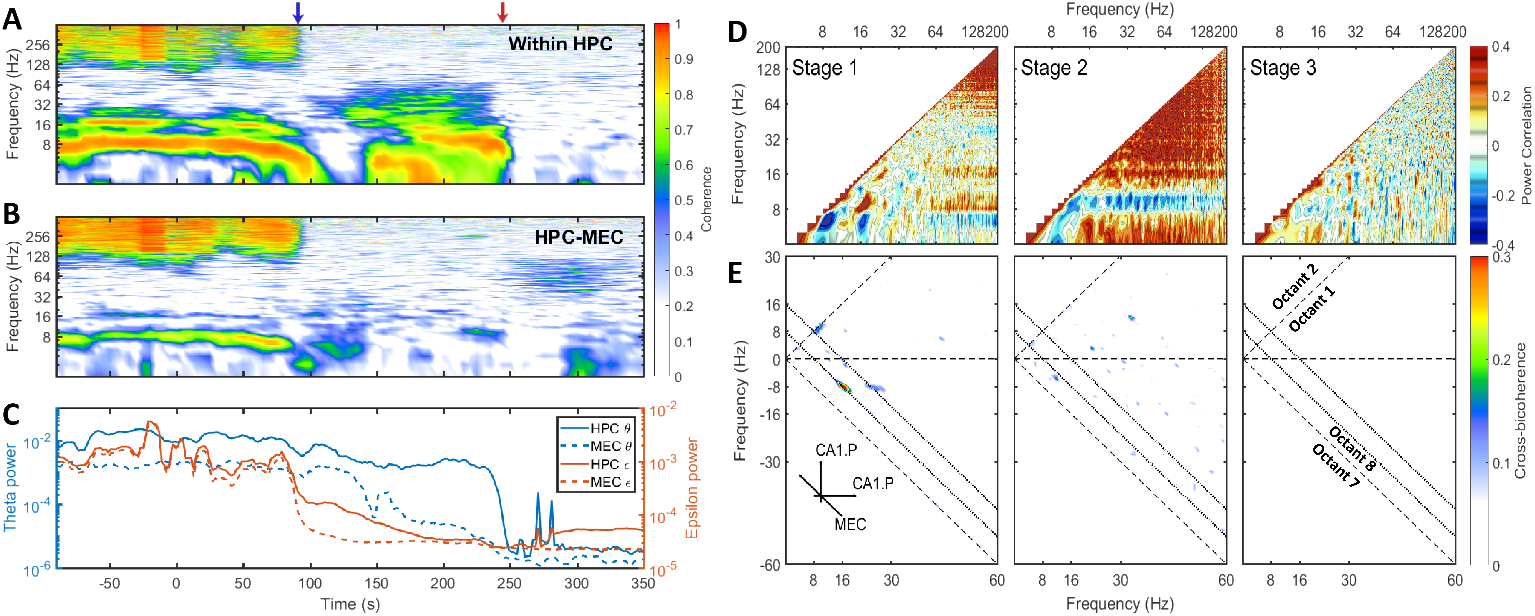
A) The development of coherence between CA1.Pyr and LM in HPC. The x-axis was the time and the y-axis was the frequency. Two transition periods were marked with blue and red arrows in the plot. Coherence is the modulus of the normalized crossspectrum with a value between 0 and 1. High coherence indicates there is a consistent phase difference between two time series at that frequency. In terms of spectral analysis of LFP, high coherence within a narrow frequency band is a sign of the existence of oscillatory rhythm, and there is a phase difference because the rhythm is a traveling wave or due to the dipolar nature of neurons. B) The development of coherence between HPC and MEC. C) The energy evolution in theta band and >100 Hz band in HPC and MEC, the variances were calculated with a window length of 10 seconds and a step increment of 2 seconds. D) Average auto-correlation coefficients of Fourier transform estimated at three stages. As the auto-correlations are symmetric, only one half was presented. Positive correlation indicates the power of those two frequency components tend to grow or decay simultaneously, while negative correlation demonstrates that the power in some frequencies is lost as others increase (see Masimore et al., 2004, 2005). During stage 1, the positive correlation can be identified in three regions: The correlation between theta and theta harmonics represented by significant dots with frequency under 32 Hz; The correlation between >100 Hz and theta (or theta harmonics) represented by horizontal strips; The correlation in >100 Hz region. During stage 2, the power of 10 Hz frequency components was negatively correlated with other frequency components, while all other frequency components had positive power correlation. E) Cross-bicoherence between HPC and MEC. Each point in the plot represented the crossfrequency coupling of the frequency triad *(f_x′_f_y′_f_x_* + *f_y_*) where *f_x_* and *f_y_* were frequency components in HPC and *f_x_ +f_y_* was frequency component in MEC. Octant 1 and Octant 8 contained all the non-redundant information. Two inclined dotted lines indicated 8 Hz and 16 Hz in MEC. During the stage 1, there were significant regions representing cross-region cross-frequency coupling of theta and its harmonics between HPC and MEC ((8,8,16) Hz, (16,—8,8) Hz and (24,—8,16)Hz). Data from r782

First, a linear and nonlinear spectral analysis was conducted to investigate the coupling in theta band within HPC and across regions. During stage 1, strong coherence existed at theta and theta harmonics frequency range across HPC layers (figure 3.5 A). The high coherence was consistent with the observation that the hippocampal LFP was dominated by nonlinear theta waves, and the theta oscillation experienced a phase reversal at hippocampal fissure [Winson, 1974]. The coherence at theta rhythm can also be observed between HPC and MEC regions, which was expected given the strong reciprocal connections between these structures and the observation of traveling theta waves in both regions [Lubenov and Siapas, 2009, Hernández-Pérez et al., 2020]. During stage 1, aside from high coherence at theta, there were significant coherent area at >100 Hz range. High coherence at >100 Hz range was not caused by oscillatory waves, but the highly synchronized >100 Hz bursts (that is, burst happened to be generated locally, but occur synchronously; figure 3.5, figure 5.3, figure 5.4 panel A&B). To illustrate the >100 Hz bursts were not caused by mechanical noise but were correlated with power of theta rhythm, a power correlation analysis [Masimore et al., 2004, 2005] was conducted, showing that during stage 1, the power of theta, the power of theta harmonics, and the power in >100 Hz range were all positive correlated (figure 3.5 D). In terms of nonlinear cross-frequency coupling. the nonlinearity of theta can be investigated through the use of the bispectrum, expressed as a significant region at the frequency triad (8, 8,16) Hz (figure 3.1). The cross-frequency coupling existed not only within hippocampal region, but also between HPC and MEC regions. Figure 3.5 E showed the cross-region cross frequency coupling for the frequency triad (*f_x_,f_y_,f_x_ +f_y_*) where *f_x_,f_y_* belonged to CA1.Pyr and *f_x+y_* belonged to MEC. revealed that theta and theta harmonics were cross-frequency coupled across HPC and MEC regions during stage 1. To summarize, theta rhythm in stage 1 had following features: 1) The oscillations at theta range were correlated within HPC, and between HPC and MEC. 2) When the power of theta was strong, high order theta harmonics would occur due to nonlinear interactions. 3) The occurrence of intermittent > 100 Hz bursts was correlated to the power of theta and theta harmonics.

After stage 1, >100 Hz bursts vanished according to the rapid power and coherence decay in corresponding band. However, there was considerable power persisting in the 6-10 Hz frequency band at the end of stage 2. Specifically, there were high coherence value around 10 Hz within hippocampus (figure 3.5 A). However, the across region coherence was weak (figure 3.5 B). Moreover, as the majority of frequencies degraded together, the power correlation during the second stage-except for the 10 Hz oscillation-were positive. That was consistent with the “parallel” spectrum degradation where all the frequency component experienced power decay. The 10 Hz oscillation, however, had negative power correlation because they experienced a power increase at the end of stage 2. The cross-bicoherence also showed that the cross-frequency coupling between HPC and MEC was weak during stage 2 compared with stage 1 (figure 3.5 E). Although the oscillation had a frequency close to theta, and high coherence within HPC region, it was weakly correlated between HPC and MEC and it was not correlated with other frequency bands. Therefore, it is most-likely distinct from theta as it does not engage a large population of neurons across brain regions, but is perhaps related to a local hippocampal network dynamic (e.g, O’Keefe and Recce, 1993).

### 3.3. Uniform Exponential Power Decay at the Second Stage

In the previous section we have shown that during stage 2, except for the 10 Hz oscillation, the power of all the frequency components were positively correlated in that their power receded together. In this section, we investigated how power of different frequency components evolved during the entire euthanasia process (figure 3.6 A). According to the power evolution plot there are two periods of rapid change: 1) From stage 1 to stage 2, high frequency components experienced rapid decay due to the disappearance of transient bursts (blue arrow and dashed line in figure 3.6 A). 2) From stage 2 to stage 3, low frequency components experienced another rapid decay because of the collapse of power-law spectrum (red arrow and dashed line in figure 3.6 A). Between these two transition periods, the power evolution of frequency components from 20 to 200 Hz can be approximated as straight lines. In the semi-log plot the straight line evolution can be interpreted as exponential decay (stage 2 indicated as red box in figure 3.6 A). The slopes of these power lines reflected the decay rates of corresponding frequency components, and during stage 2 the power evolution were almost parallel which indicated that frequency components shared a similar decaying rate.

**Figure 3.6.**
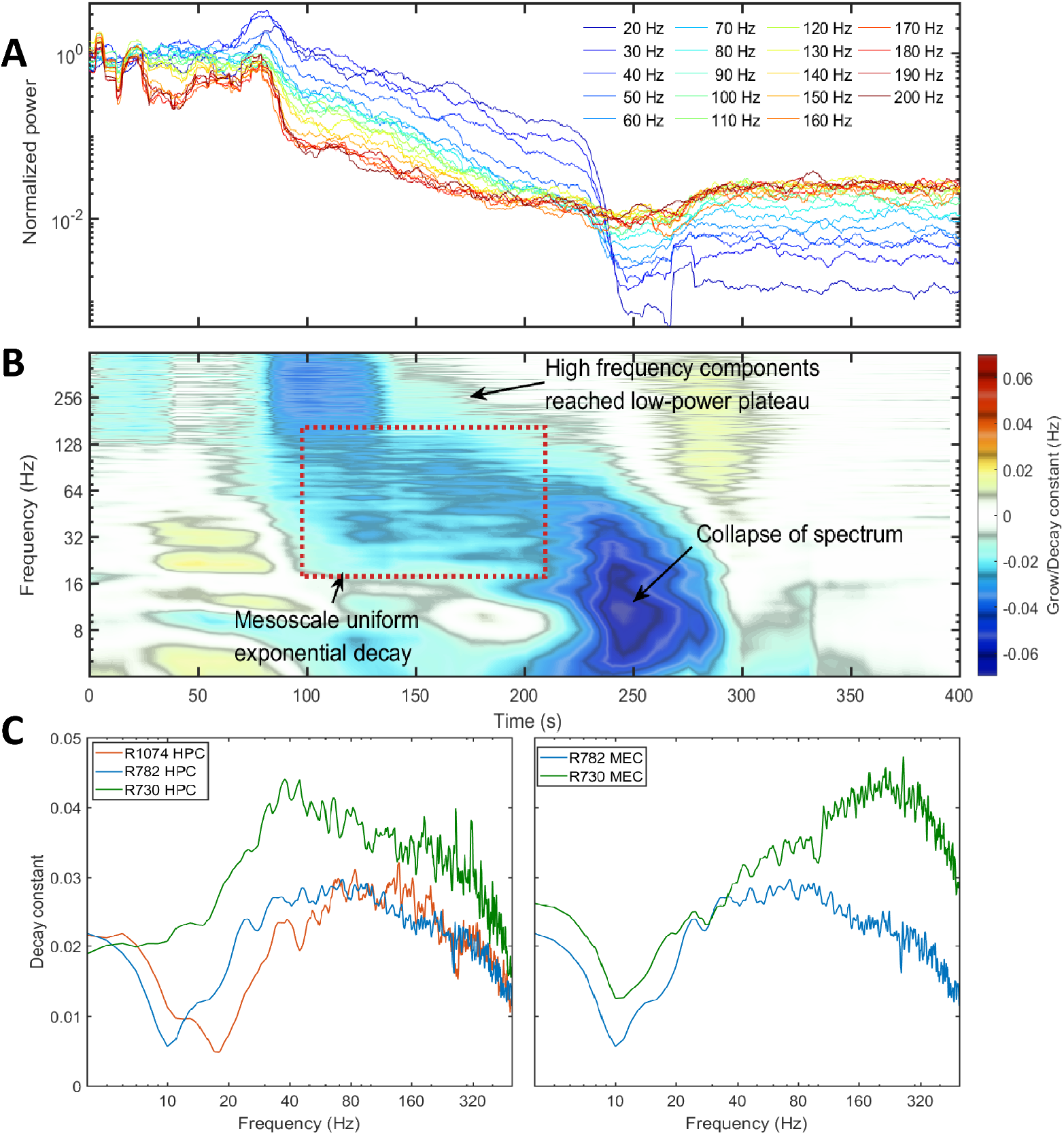
Power decay of frequency components over the euthanasia process. A) The power evolution for frequency component from 20 Hz to 200 Hz. Each line corresponded to the power decay of one frequency component. Low frequency components were indicated by cold colors and high frequency components by warm colors. Note the the power were normalized by their value at the time of injection (*t* = 0). Data from rat 782. B) The percentage power change rates of different frequency components over the entire spectrum degradation process. To obtain the percentage power change rate, the power time series for each frequency component was first estimated and the percentage power change rate was defined as the ratio of the time derivative of power time series to the power time series. If a frequency component experiences a exponential decay or grow *e^αt^*, the percentage power change rate will be a constant with value *α*. Data from rat 782. C) The decay constant versus the frequency during the exponential decay stage. The left panel were the decay constants in HPC among rats and the right panel were decay constants in MEC. The decay constant were estimated by averaging over the entire stage 2 period.

The decaying rate (in the unit of Hz) can be quantified by differentiating power time-series with respect to time, and normalizing the time derivative by the power (figure 3.6 B). At the transition period from stage 1 to stage 2, both theta and high frequency component expressed a decay in power. During stage 2 of spectrum degradation, the frequency components over 20 Hz decayed at similar rate with a decay constant around 0.03 Hz. This exponential decay of high frequency components lasted shorter as their power reached the low energy plateau and experienced limited power decay afterwards. Stage 2 ended with a rapid decay of low frequency components with decaying rat of 0.06 Hz corresponding to the collapse of the power-law spectrum.

The stage at which power decayed exponentially with similar rate of decay was observed across animals and regions (figure 3.6 C). The decay rates of frequency components lower than 20 Hz were small due to the existence of 10 Hz rhythm at the end of stage 2. The spectrogram in figure 3.2 indicated that the frequency of that oscillation changed from around 20 Hz to 10 Hz during the degradation process and became more prominent compared with power of frequency components at adjacent region. In the spectra evolution (figure 3.2, figure 5.1 & figure 5.4 A) the 10 to 20 Hz oscillation acted as a spectrum front, but the reason of its generation and development is unknown. Apart from that, all the frequency components had a exponential decay with decay constant around 0.02 in HPC and 0.04 in MEC. High frequency components had relatively slower averaged decay rate due to the existence of low power plateau.

## 4. Discussion

The current manuscript investigated the degradation of the hippocampal power spectral density over the course of euthanasia. Specifically, while the slope of the spectra has been known to change as a function of running speed [Sheremet et al., 2019b], the parametric space of the 1/f slope is small relative to specific bands such as theta. Given that it exists across the spectrum from the “low activity” sleep states to instances when the hippocampus receives a large amount of synaptic input suggests that it is a persistent baseline state. However, the mechanisms that support the 1/f slope and the degree to what is “true” activity is unknown.

Therefore, euthanasia offers the opportunity to observe the spectral collapse as the amount of synaptic activity wanes.

We observed three states of spectral degradation. Stage 1, or pre-effective stage, can be described as a typical “active”, theta-dominate state. In stage 2, or quasi-stationary decay stage, the peak above 1/f in the theta range was eroded as was the higher frequency, >100 Hz bursting. Throughout this stage, the power in the higher frequencies of the power spectra eroded with higher frequencies such as 120 Hz reaching the absolute electrothermal noise background before lower frequencies (such as 70 Hz). Interestingly, this collapse in the higher frequencies (including the 60-120 Hz gamma band) was associated with the appearance of a 10 Hz peak in the spectra in the hippocampus. The duration of stage 2 persisted longer in the hippocampus relative to the medial entorhinal cortex (which did not exhibit a 10 Hz peak). The end of stage 2 and start of stage 3, or collapse stage, was marked by a complete collapse of the power in all spectral bands, with low activity in nearly all bands and occasional large spikes that plausibly related to spreading depolarization [Pani et al., 2018]. Finally, the activity reached a minimum, related more to the electro-thermal noise in physical recording environment surrounding the data acquisition system than neural activity.

In light of this, we interpret the 1/f spectral degradation during euthanasia to be in line with the energy cascade model. Specifically, rhythmic activity is spontaneous generated by the reentrant architecture in the brain [Edelman, 1987, Tahvildari et al., 2012, Edelman and Gally, 2013]. The frequencies of rhythmic activities cover a wide frequency range because the oscillators have different intrinsic frequencies depending on the size of reentrant architectures. Low frequency, high power rhythms are a function of large scale activity that extends beyond a single brain region whereas high-frequency, low amplitude oscillations are local and supported by interactions within a specific brain region [Buzsaki and Draguhn, 2004, Buzsaki, 2006, Sheremet et al., 2019a].

Previously, we have argued that the organization with a spectral band is a function of the activity within the hippocampus [Zhou et al., 2019]. Briefly, during normal behavior, as animals increase their running speed, the amplitude of theta and its harmonics increase. The increase in these frequency bands is accompanied by a loss of power in neighboring bands (e.g., 5-7 Hz; 10-14 Hz). With more excitatory input into the network, neurons are more capable of pushing an pulling on each other. If a neuron or a group of neurons is oscillating faster than the plurality, they are “slowed down”. If neurons are oscillating too slow, the population en masse, speed them up. A “collective enhancement of precision” is achieved [Strogatz, 1994] where one frequency band increases in power at the expense of neighboring bands.

Therefore, if theta power is a consequence of activity dependent entrainment, then we hypothesize that the global loss of theta during stage 2 represents an inability of neurons to entrain each other due to the loss of energy. This is supported by the synchronous erosion of visually detectable theta across regions (e.g., MEC and HPC), with this activity become dispersed and slowed into the 1-4 Hz range. Lower levels of activity are still moving through the larger reentrant loops, but the activity is no longer coordinated. Stated differently, the physical architecture, along with its synaptic time constant and axonal conduction delay still exists. Moreover, activity is still moving through this larger reentrant loop supporting the power in the lower frequency bands. However, the activity is simply uncoordinated, with some pulses moving at 6 Hz and others at 9 Hz. Without the coordinated precision, there is no clear peak in the spectra (n.b., from this perspective, it is imprecise to suggest that an oscillation is missing or absent. Rather, the activity is simply not cohesive). However, the activity is still sufficient to drive the smaller, local circuits supporting higher frequency rhythms. For instance, at the start of stage 2, there was significant power in the gamma range that gradually eroded with the higher frequencies merging with the electro-thermal background first.

As the hippocampus is a densely recurrent network of neurons [Lorente de No, 1938], then barrages of excitatory input from the large scale will drive the smaller circuits and cast higher frequency activity, such as gamma (see figure 1c of Berg et al., 2019). The frequency of gamma has been theorized to be proportional to the amount of excitatory input; more input, higher frequency [Traub et al., 1996, Ahmed and Mehta, 2012, Zhou et al., 2019]. Therefore, as the activity in the large scale loops become less coordinated and reduces in strength, the higher frequency range of gamma will succumb first. Interestingly, the time course of the gamma erosion is approximately three times longer in the hippocampus relative to the medial entorhinal cortex, which may be due to either the higher amount of reciprocal activity of neurons within the hippocampus, higher excitability of this region during a loss of perfusion, the higher degree of connectivity of the hippocampus with other brain regions, or a combination of these. Perhaps more interesting is that towards the end of stage 2, the erosion of gamma power appears to stop at ~10 Hz in the hippocampus, which may represent the intrinsic membrane resonance properties of the pyramidal cells and a subpopulation of interneurons [O’Keefe and Recce, 1993, Kamondi et al., 1998, Yamaguchi and McNaughton, 1998, Bose et al., 2000, Magee, 2001, Lengyel et al., 2003, Maurer et al., 2005, 2006, Geisler et al., 2007]. When the excitatory input into the hippocampus declines to the point that it is no longer able to support the lower 10 Hz oscillation, there is a complete collapse of the spectrum to quiescent levels.

Up to this point, we have defended the idea that the 1/f spectra is supported by the cascade of activity from large reentrant circuits into smaller, nested loops [Buzsaki and Draguhn, 2004, Buzsaki, 2006, Sheremet et al., 2019a]. An alternative hypothesis exists in which the peaks above the 1/f slope are true oscillators, where as the slope itself is the consequence of a distinct broadband, arrhythmic activity [He, 2014] or similarly, rhythmicity superimposed onto a wide-band, noisy background [Bullock et al., 2003]. To argue divisions between the periodicity versus aperiodicity in the LFP based on the Fourier power spectral density is to risks misinterpreting the decomposition for an interpretation and potentially obfuscates the biophysics of the system.

First, in terms of the decomposition, it should be appreciated that while the “one amplitude to one frequency” assignment of the power spectral density give the impression that each frequency is independent of each other, they most certainly are not [Bullock et al., 1995, Buzsaki, 2006]. The same synapses and neurons contribute to all rhythms simultaneously. As a simple example, theta casts harmonics that can go upwards of 56 Hz [Sheremet et al., 2020]. A single event can appear in multiple locations of the power spectra. More nuanced is that the same synaptic events that support theta also support gamma. The post-synaptic potentials, carrying the majority of the LFP, can be entrained to both theta, the harmonic of the theta and gamma. It is the decimation into sinusoids that allows one to describe “three components”, which are in fact one and the same. To extend this one step further, as synaptic events do not respect the sinusoidal decimation, the manner in which the LFP is decomposed allows a single event to contribute to “arrhythmic” and “rhythmic” bands simultaneously. This concept has been illustrated before in which the time-series that makes a Bach concerto can be played in reverse or chopped and reordered to the point in which the most ardent scholar no longer recognizes it and yet, the power spectra remains unchanged [Buzsaki, 2006]. A musical trace can be robbed of the rhythmicity that allows one to identify it. However, if the power spectral density arrives at the same realization, then it does not make sense to use it for a “periodic”/”aperiodic” division. This concept has also been demonstrated previously in which synthetic “arrhythmic” and “rhythmic” time series can give way to identical power spectral density profiles (Figure 4 of Sheremet et al., 2016). It needs to be appreciated that the decimation into a sum of sinusoids is arbitrary, dividing non-sinusoidal rhythms or cross-frequency dependent events into an artificial and potentially misleading representation if the underlying physiology is not well understood.

Second, the division into rhythmic and arhythmic components falls short when considering the biophysics of the system. The rationale for ascribing “arhythmic noise” to the background was perhaps due to the unintentional representation of independence among frequencies in the power spectral density representation (see above). Specifically, it was suggested that individual oscillators aligned to each frequency would be more magic that biological reality [He, 2014]. As noted above, a synaptic event can contribute to power in multiple rhythmic and arrhythmic bands simultaneously. Therefore, there does not need to be an alignment of oscillators at each individual frequency. In fact, noise has been defined as the ratio between excitation and inhibition [Voytek et al., 2015], which is the same process that support the “rhythmic activity” [Berg et al., 2019]. Therefore, while it may be tempting to dissect to power spectra into periodic and aperiodic components [Donoghue et al., 2020], as a single synaptic event can contribute to signal and noise simultaneously, the tool itself has no insight as to whether the underlying system is rhythmic or not. Moreover, peak over the 1/f slope has been argued to be incomplete in defining an oscillation elsewhere [Pesaran et al., 2018]. (Rosen, 1991, p. 60): … There can be no greater act of abstraction than the collapsing of a [natural] phenomenon down to a single number, the result of a single measurement. From this standpoint, it is ironic indeed that a mere observer regards oneself as being in direct contact with reality and that it is ‘theoretical science’ alone that deals with abstractions.

This sort of either/or classification has been endemic since nearly the start of the field (perhaps unintentionally stemming from the reductionist approach that dominated the field at the time) in which terms were applied to flutters of a certain class. Researchers concocted the terms “theta”, “gamma” and “noise” giving them individuality and independence when, in fact, neurobiology makes no such distinction – and they themselves are identical. It is worth recovering that the bands with a name in fact describe dynamics at a scale. Theta is the propagation of activity along a large “macroscale” loop. Nested within in this large loop are many smaller loops in which the activity can be reciprocally coordinated on the mesoscale, supporting faster rhythms such as gamma [Buzsaki and Draguhn, 2004, Buzsaki, 2006]. The scales – and the names that define them- are not orthogonal but intricately coupled.

Of course, the question remains, what is responsible for the 1/f sloped background when there are a finite number of reciprocal loops in the nervous system? The answer to this question is best addressed through an analogy, using a system defined by forcing and nested loops of multiple scales – the cardiovascular system. Should one measure the velocity of a red blood cell in a capillary, the time-series itself will exhibit the dominant frequency of the heartbeat (macroscale forcing event). As heartbeats are “slow charge, rapid discharge” event, the large amplitude changes in velocity will exhibit a significant deviation from a sinusoid. Furthermore, this forcing occurs as a cascade from macro to micro scale, from arteries through arterioles to capillaries. Thus, blood cell velocity will be subjected to other influences, such as friction as a result of running into the other blood cells or the vascular walls or even form high frequency turbulent eddies (e.g., partial occlusion). This jostling can be identified in the velocity profile as being a repeatable, low amplitude-high frequency event coupled to the macroscale heartbeat frequency. Finally, decomposing the red blood cell velocity would reveal a power spectra density eerily similar to the hippocampus, complete with a fundamental frequency, harmonics and a 1/f background (see figure 2B of Harlepp et al., 2017). In fact, one may suspect that the influence of pentobarbital on the power spectra of red blood cell velocity would parallel the degradation observed in the LFP, with the highest frequencies succumbing first. Therefore, while Fourier decomposition is certainly a useful tool in time-series analysis, defining the presence of an oscillation as a peak above the background and the absence as having no peak along the 1/f background as “aperiodic” from the power spectra has a peculiar and contracted relevance. The cardiovascular system never evolved to make rhythms or arrhythmic activity, but to provide circulation which comes in the forms of beats. While the study of the rhythms may be diagnostic, providing insight into how well the cardiovascular system is functioning, separating rhythm from nonrhythm based on the power spectra provides limited insight into how the red blood cell moves.

As “...the EEG reflects the “average” behavior of neurons” (Buzsaki, 2006, p. 129), this analogy holds for the nervous system. Neurons are the microscale components that reside in nest loops of multiple scales. The LFP is the aggregate activity related to activity moving through multiple loops of different scales simultaneously. The movement of the activity through the nervous system is, from this perspective, a unitary process where activity chases is own activity through reentrant loops [Hebb, 1958]. Certainly, there may be activity that expresses as a peak above the 1/f background, such as theta which comprises the movement of activity on the macroscale. However, oscillations like gamma rarely peak above the background [Zhou et al., 2019] and yet also describe the mesoscale volleys of activity governed by interneurons. Therefore, appreciating the power spectra as a decomposition of activity moving from the macro- to microscale, then it is a false dichotomy in making a “non-rhythm/rhythm”, “present/absent”, “periodic/aperiodic” distinctions. Cross-frequency dependence is the rule rather than the exception.

## 5. APPENDIX

**Figure 5.1.**
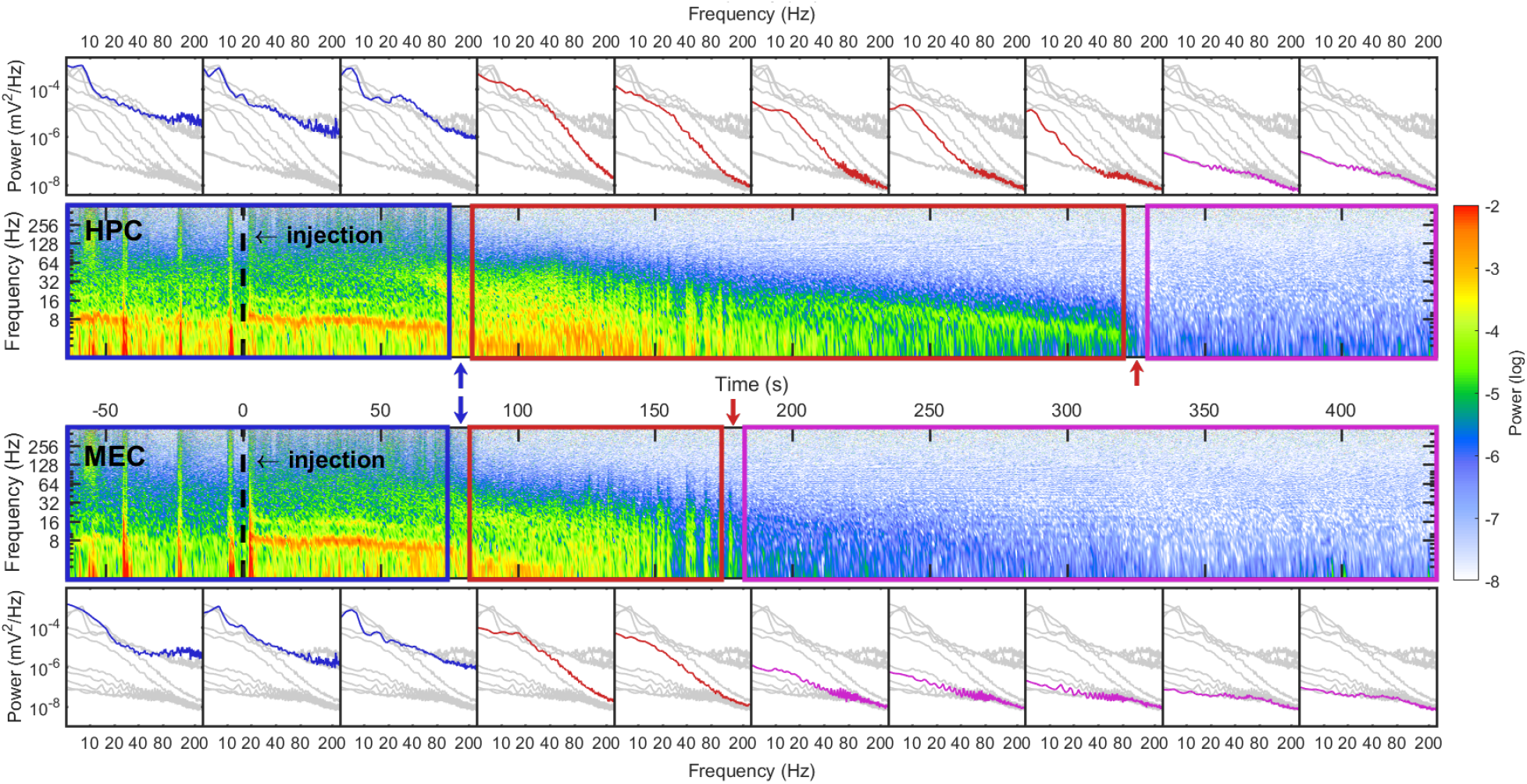
Similar with figure 3.2, Data from rat 730.

**Figure 5.2.**
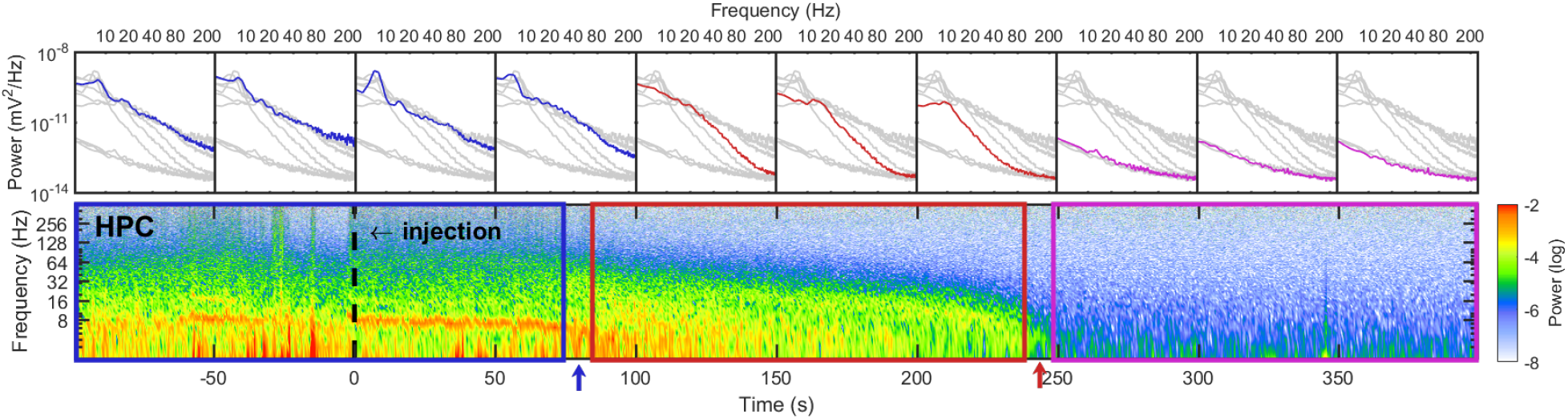
Similar with figure 3.2, spectrum degradation in hippocampus (HPC).

**Figure 5.3.**
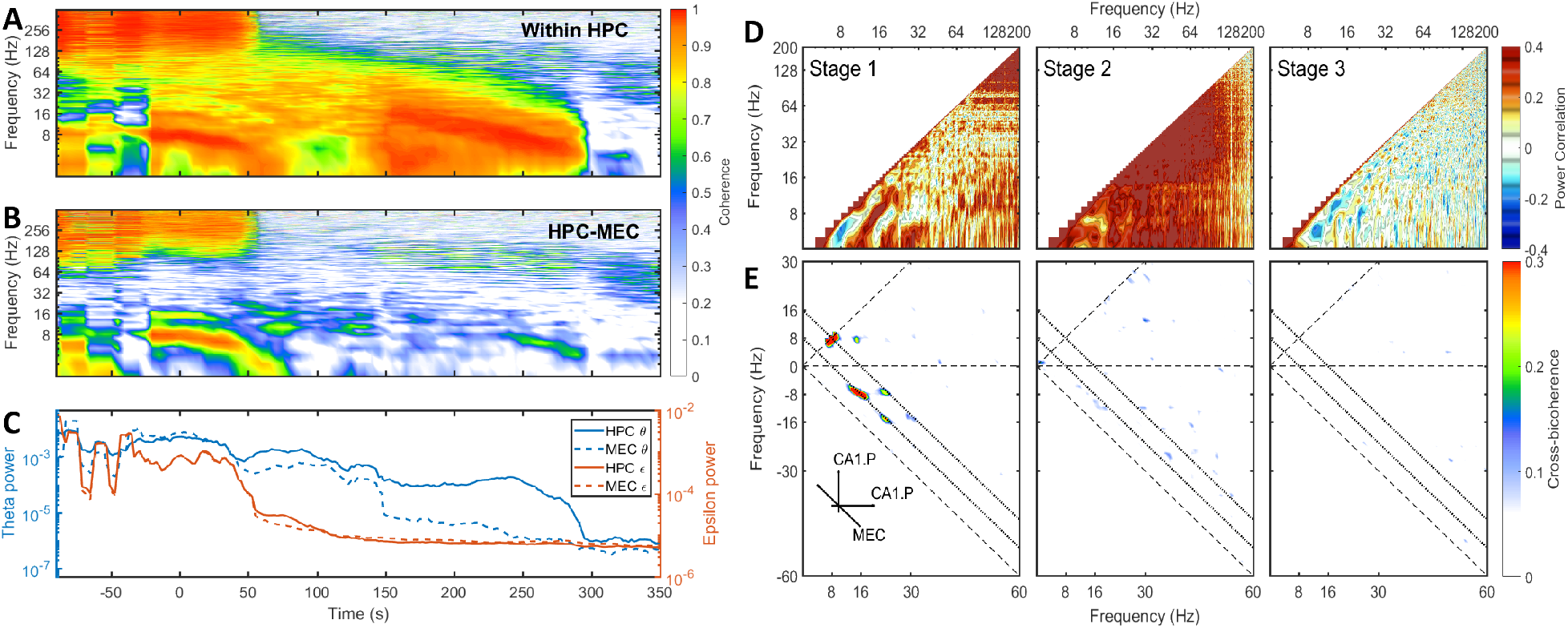
Similar with figure 3.5. Data from rat 730.

**Figure 5.4.**
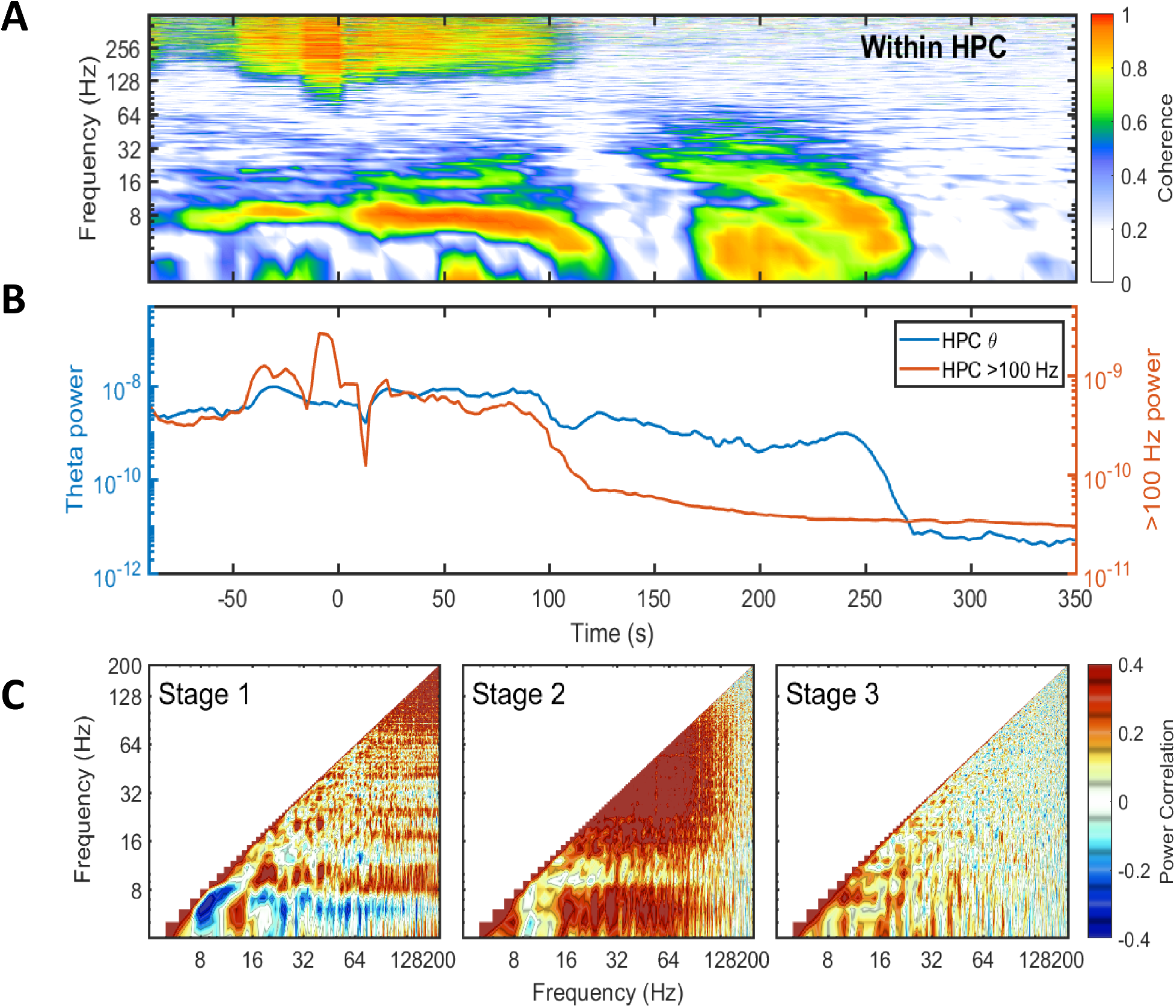
Similar with figure 3.5 without cross region coherence and bicoherence Data from rat 1074.

## Notes

Funding information This work was supported by the McKnight Brain Research Foundation, and NIH grants. Grant Sponsor: National Institute on Aging; Grant number: AG055544 and Grant Sponsor: National Institute of Mental Health; Grant Number: MH109548.

### Competing Interest Statement

The authors have declared no competing interest.

