## Supplementary figures and images for "Spectrum Degradation of Hippocampal LFP During Euthanasia"

### Supplemental Figure 1

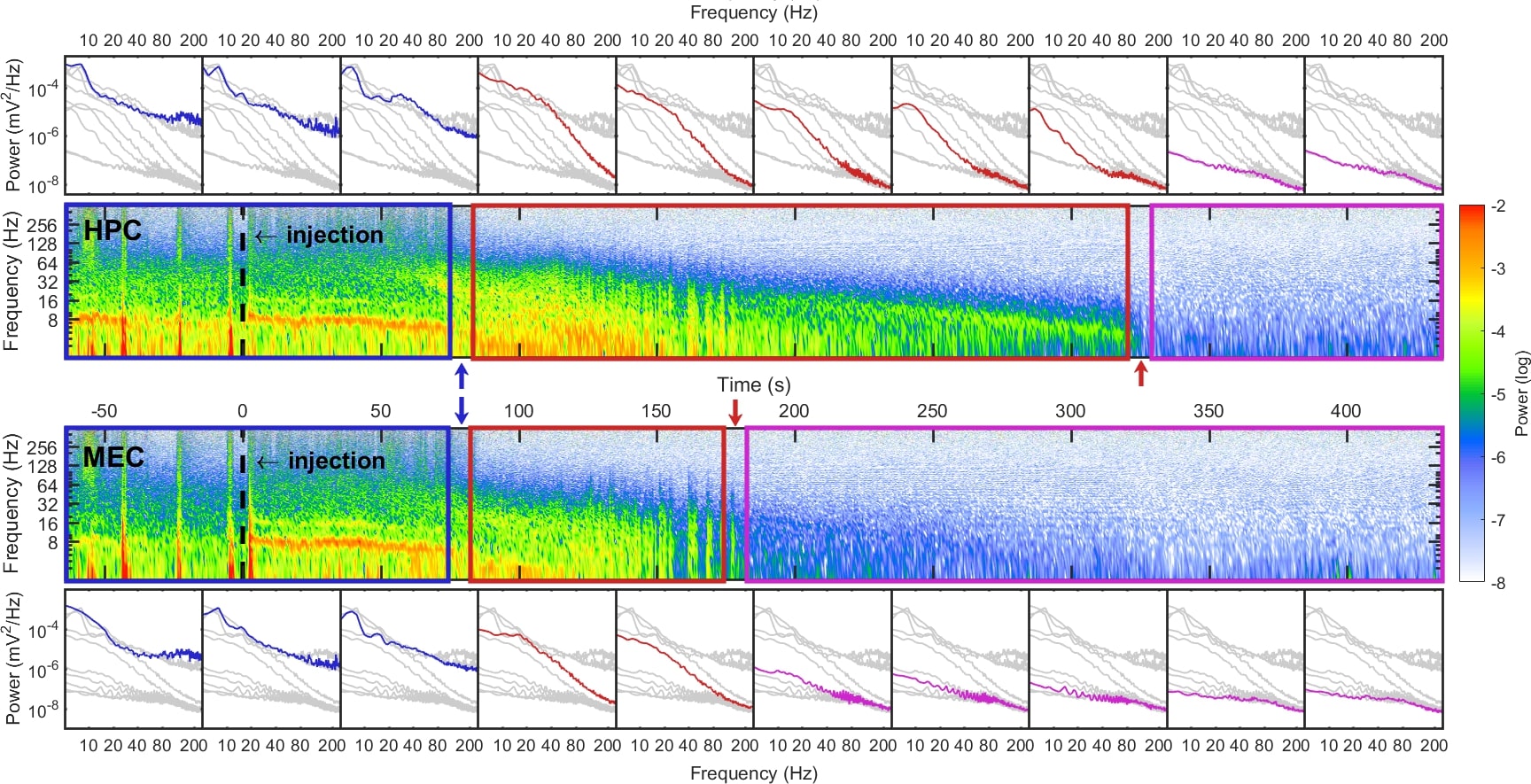

### Supplemental Figure 2

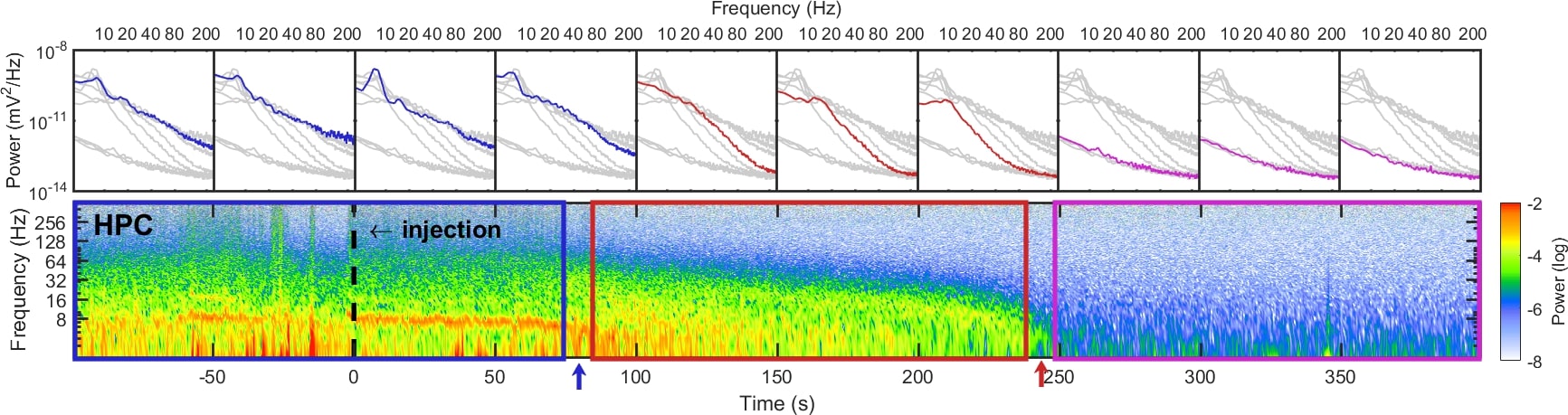

### Supplemental Figure 3

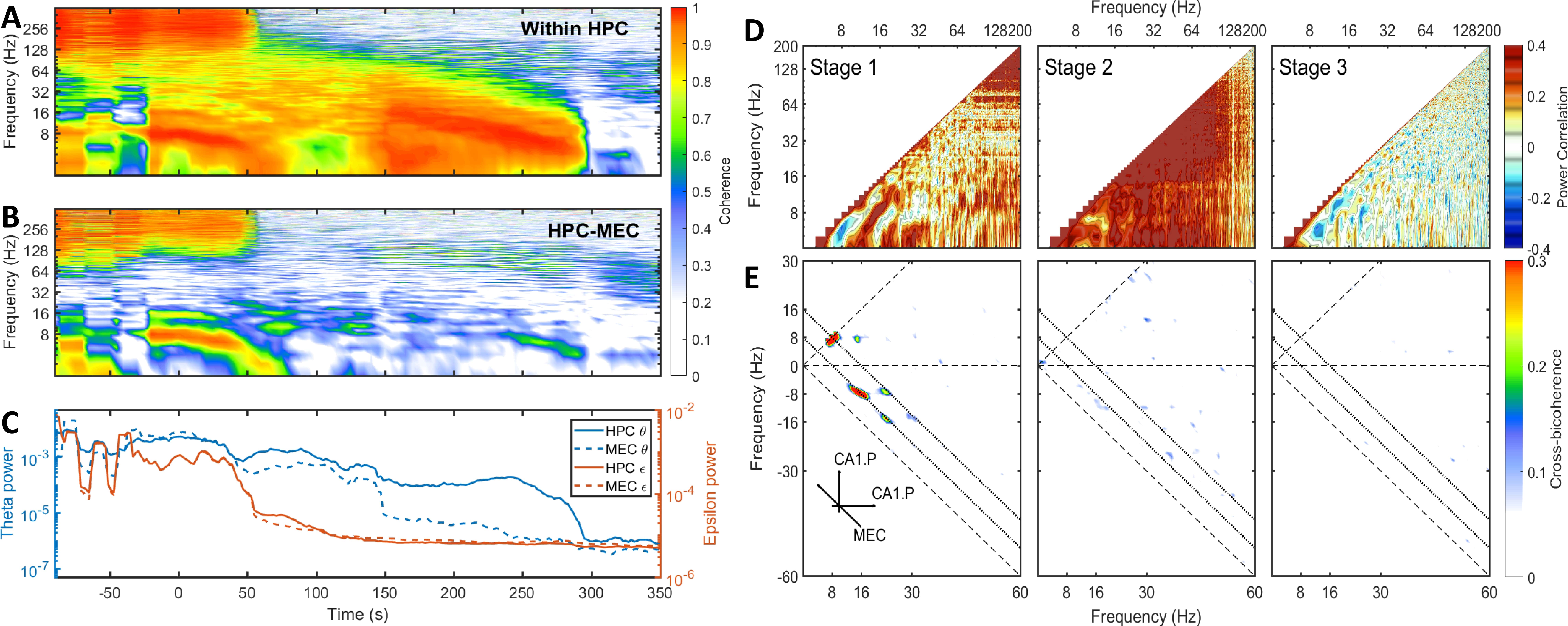

### Supplemental Figure 4

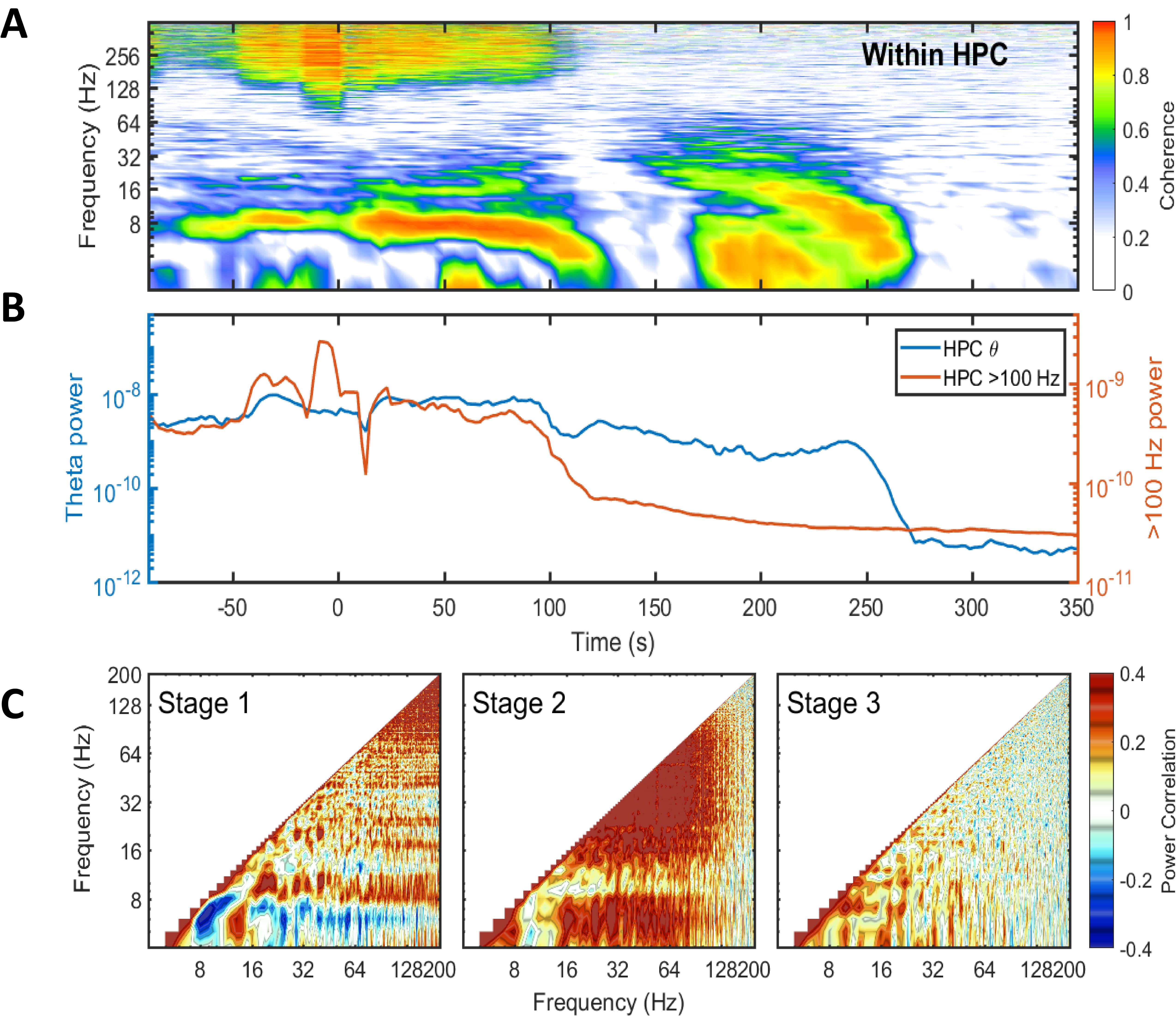
